# 3D-Printed Titanium Implants with Bioactive Peptide-polysaccharide Scaffolds for Personalized Bone Reconstruction

**DOI:** 10.1101/2025.09.08.674817

**Authors:** Noam Rattner, Vladimir Perlis, Eran Golden, Ariel Pokhojaev, Rachel Sarig, Itzhak Binderman, Michal Halperin-Sternfeld, Solomon Dadia, Lihi Adler-Abramovich

## Abstract

Large bone defects caused by trauma, tumor resection, or congenital abnormalities remain a major clinical challenge. Standard titanium implants are widely used due to their strength and biocompatibility, but their bioinert surfaces often lead to poor osseointegration. The emergence of 3D printing has enabled patient-specific titanium implants with tailored architecture and mechanical properties. However, these constructs still lack the bioactivity required for robust and spatially uniform bone integration, particularly within the implant core. To address this limitation, we developed a bioactive, cell-free strategy that integrates porous titanium implants with a nanofibrillar peptide-hyaluronic acid scaffold, delivered either as a hydrogel or in lyophilized form. The scaffold exhibited enhanced enzymatic stability and supported osteoblast-like cell adhesion *in vitro*. In a rabbit calvarial critical-size bone defect model, scaffold-integrated implants significantly outperformed inert controls, with hydrogel integration nearly doubling inner bone volume and improving trabecular architecture. Histological analysis confirmed enhanced bone-implant integration, active periosteum, healthy marrow, and reduced inflammation. This acellular, growth-factor-free approach combines the structural precision of titanium with the regenerative potential of ECM-mimicking scaffolds, offering a translatable pathway for personalized skeletal repair.

## 1. Introduction

Bone defects caused by trauma, tumor resection, or failed surgical procedures frequently affect load-bearing bones, where structural integrity and long-term mechanical performance are critical^1,2^. When such defects exceed the body’s natural regenerative capacity, they are considered critical-size defects and represent a major clinical challenge^3–5^. Although bone is a dynamic tissue capable of self-repair when fracture fragments are stabilized and closely aligned^6,7^, regeneration in critical-size defects is impaired due to disrupted biological and mechanical environments^7,8^.

To restore both structure and function in such cases, surgical intervention is required, often including the implantation of a device that provides stability or replaces the missing bone^1,9,10^. Traditional approaches such as autografts, allografts, and standard prosthetics can fall short, especially in anatomically complex cases, highlighting the need for more effective solutions^3,5^.

A widely adopted solution is the implantation of metallic devices, particularly those composed of titanium or titanium alloys, which are favored for their strength, biocompatibility, and corrosion resistance^11–13^. The emergence of additive manufacturing has enabled the fabrication of patient-specific, three-dimensional (3D) printed titanium implants offering tailored geometry and mechanical properties^4,9,10,14–16^. In some cases, the use of patient-specific implants in bone defects adjacent to a joint can serve as an alternative to limb amputation^17^. However, the bioinert nature of titanium limits direct interaction with host tissue, often resulting in poor osseointegration and limited new bone formation within the inner porous regions of the implant. This lack of bone ingrowth into the core often leaves large central voids, weakening the implant’s stability and putting the long-term success of the reconstruction at risk, especially when the implant has to bear load^8,14–16,18–20^.

To address these limitations, numerous strategies have been developed to enhance the bioactivity of porous titanium implants. One major approach has focused on surface modifications of the printed structure. Techniques such as acid etching, alkali treatment, anodization, and plasma spraying have been employed to tailor surface roughness and chemistry, thereby promoting protein adsorption, osteoblast adhesion, and subsequent bone formation^21^. In addition, biofunctional coatings, including calcium phosphate, hydroxyapatite, bioactive glasses, and peptide-functionalized layers, have been applied onto porous titanium to accelerate osseointegration and provide osteoconductive cues^21–26^. Beyond surface modifications, another strategy involves incorporating bone graft materials, such as autografts and allografts, into the porous implant structure to provide a direct osteogenic stimulus^27–29^. While these grafts possess inherent osteoinductive and osteoconductive properties, their limited availability, donor site morbidity, and risk of immune rejection make them a challenging clinical option^30,31^.

Additional efforts have turned to synthetic bone substitutes and hydrogels, which provide a more adaptable and biomimetic microenvironment. For example, silicate-substituted calcium phosphate was incorporated into a lamellar titanium cage to enhance bone formation and implant fusion in a sheep spine model^32^. Hydrogels, in particular, have gained considerable attention owing to their tunable mechanical properties, injectability, and ability to deliver cells, drugs, or bioactive molecules^33,34^. Several systems have been investigated, including hydroxyapatite-loaded calcium alginate hydrogels^35^, and silver nanoparticle-loaded hydrogels^36^. PLGA-PEG-PLGA gels have primarily served as drug carriers for cisplatin or simvastatin within the titanium implants post tumor resection, where they suppressed local tumor recurrence and simultaneously promoted bone regeneration^37,38^. Despite their promise, most hydrogel systems still face significant challenges, including dependence on controlled drug or growth factor release, difficulty synchronizing degradation with new tissue formation, limited and uneven infiltration throughout the porous architecture, and insufficient mechanical stability under load-bearing conditions^39–44^. These limitations emphasize the need for next-generation hydrogel formulations that combine biomimicry, mechanical resilience, and bioactivity to achieve consistent regeneration throughout the implant core^45^.

A promising approach involves mimicking the native extracellular matrix (ECM), which is composed of a 3D fibrillary network predominantly made of proteins, with collagen being the most abundant^46–48^. Self-assembling amino acids and short peptides can form nanofibrillar structures that resemble the ECM’s morphology^47,49–52^. These peptides are highly tunable in their mechanical and biochemical properties and can incorporate molecular recognition motifs such as the arginine-glycine-aspartic acid (RGD) sequence found in fibronectin, enabling enhanced cell adhesion and signaling^53,54^.

Traditionally, scaffolds designed to mimic the native ECM are fabricated either as hydrogels in a hydrated state or as dry, solid structures^1,47^. Lyophilization is commonly used to convert hydrogels into dry scaffolds by removing water while preserving the 3D fibrillary architecture^55,56^. However, the outcome of lyophilization depends on multiple factors, including freezing rate, fiber stability, and processing parameters, which can subtly alter the scaffold’s microstructure compared to the original hydrogel form^57^. As a result, although lyophilized scaffolds retain an ECM-like network, their mechanical and physical properties often differ from those of the hydrated hydrogels, despite originating from the same material^58^. Understanding these differences is crucial, as the scaffold’s physical state can influence cell behavior, tissue infiltration, and ultimately, the success of bone regeneration^55,59,60^.

We and others have previously demonstrated the potential of peptide-based nanostructures in a range of biomedical applications^47,54,61–63^. We demonstrated that Fmoc-diphenylalanine (Fmoc-FF) peptides can self-assemble into elongated β-sheet fibrillar networks that interpenetrate hyaluronic acid (HA), yielding homogeneous composite hydrogels with markedly improved mechanical rigidity and resistance to enzymatic degradation, while preserving biocompatibility and enabling the stabilization and sustained release of bioactive molecules without the need for toxic cross-linkers^64^. When integrated into polymeric scaffolds, HA-peptide composites were further shown to promote osteogenic differentiation and mineralization *in vitro*, highlighting their potential for bone tissue regeneration^54,65^. More recently, fibrous FmocFF/HA hydrogels demonstrated potent immunomodulatory properties *in vivo*, where modulation of macrophage polarization supported vascularization and significantly enhanced bone regeneration^66^.

In this study, we developed and studied a novel composite implant system combining 3D-printed porous Ti-6Al-4V implants with the self-assembling FmocFF/HA nanofibrillar scaffold, delivered as either a hydrogel or in lyophilized form. Using a critical-size calvarial defect model in rabbits, we investigated the scaffold’s ability to induce bone regeneration and how the physical state of the scaffold influences spatial bone formation within the implant. While the lyophilized scaffold was found to support bone formation mainly at the defect periphery, the hydrogel form was shown to promote substantial regeneration across both peripheral and central regions, exhibiting a uniform and volumetric pattern. This novel system offers a promising strategy for personalized bone regeneration in various pathologies.

## 2. Results and Discussion

### 2.1 Implant-Scaffold Integration and Characterization

The porous architecture of the titanium implant plays a key role in facilitating bone ingrowth and integration with the host tissue^5,43^. In this study, the gyroid lattice design was employed, offering a continuous, interconnected pore network that not only supports vascular and cellular infiltration but also provides mechanical stability and uniform stress distribution^4^. This geometry further enables efficient incorporation of bioactive scaffolds via simple infusion techniques. For this purpose, we immersed the implant in an HA solution, followed by the addition of the FmocFF peptide and brief vortexing, which induced an *in-situ* sol-gel transition within the pores. The resulting FmocFF/HA hydrogel was confined within the implant structure or could subsequently be lyophilized, thereby preserving its 3D architecture and facilitating later rehydration (Figure 1A-C). To evaluate the FmocFF/HA hydrogel, in its wet or its lyophilized form, infiltration into the internal architecture of the 3D-printed titanium mesh, we designed a cylindrical implant that could be longitudinally divided into two semi-cylinders for direct observation (Figure 1D). Cross-sectional observation revealed that the hydrogel uniformly infiltrated the entire mesh structure (Figure 1E-J).

**Figure 1.**
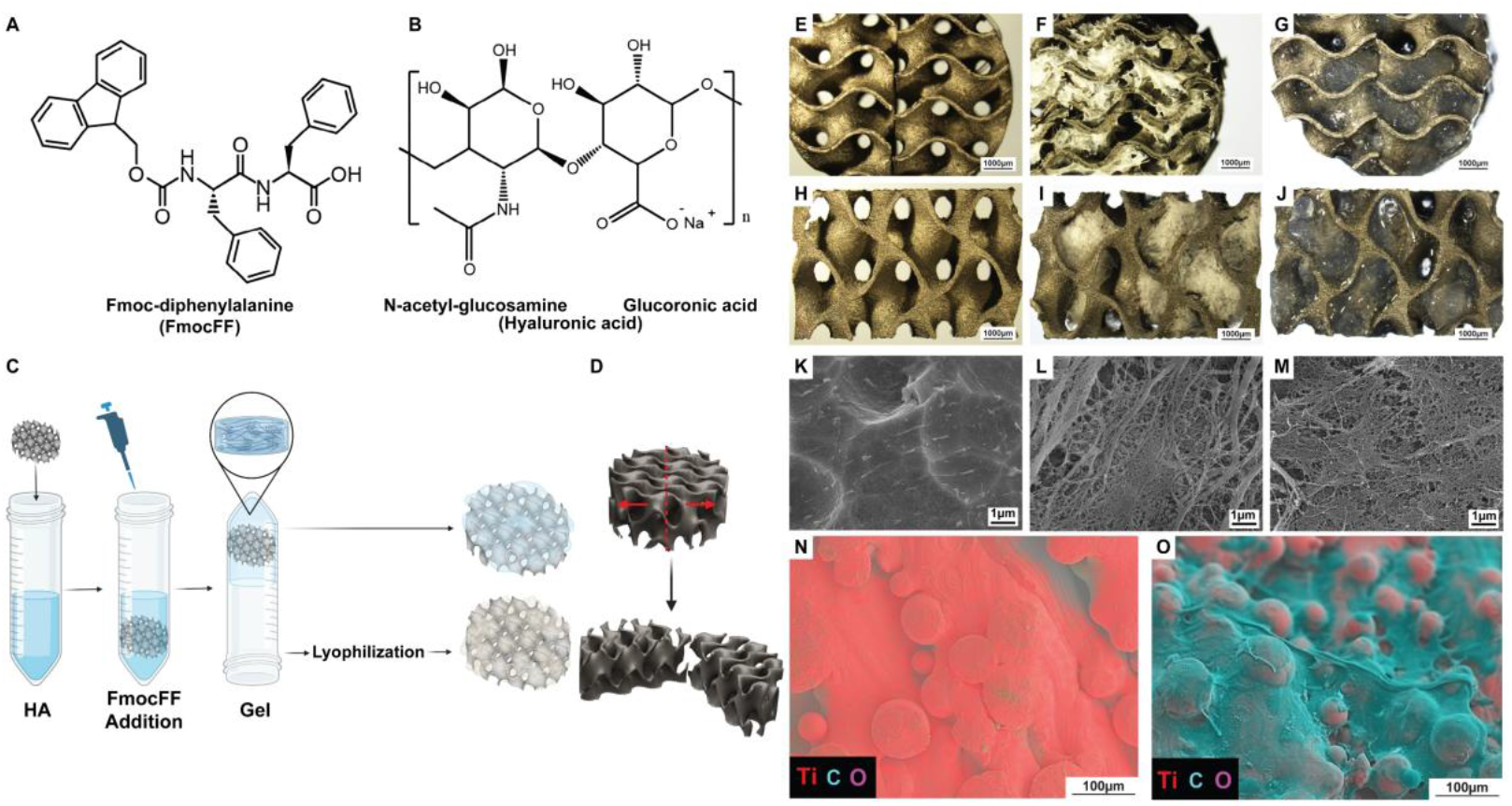
*In vitro* characterization of 3D-printed implant lattice incorporating FmocFF/HA scaffolds. (A) Molecular structure of the Fmoc-diphenylalanine peptide. (B) Molecular structure of HA. (C) Schematic illustration showing the incorporation of a hydrogel or lyophilized scaffold into the 3D-printed implant. (D) Illustration of the custom 3D-printed lattice fabricated for the *in vitro* scaffold penetration assessment. (E-G) Top-view images. (H-J) Lateral view images. (K-M) HRSEM images displaying the surface. (N, O) EDS analysis. (E, H, K) Inert implant, (F, I, L) Lyophilized FmocFF/HA-incorporated implant, (G, J, M) Hydrogel-Incorporated Implant, (N) Inert Implant, (O) Lyophilized FmocFF/HA-incorporated implant.

As an alternative integration strategy, we injected the preformed FmocFF/HA hydrogel directly into the implant using a fine cannula. Despite the hydrogel’s self-healing capabilities^66^, this approach resulted in partial leakage. The high shear stress and pressure generated within the narrow lumen of the cannula disrupted the hydrogel network, leading to a compromised scaffold integrity. Based on these observations, we selected the two more effective methods for further investigation: *in-situ* gelation within the implant pores and lyophilization following gelation.

HRSEM imaging revealed a network of fine fibrils formed by the self-assembly of the FmocFF peptide, both in the hydrogel state and after lyophilization (Figure 1K-M). These fibrillar structures closely mimic the 3D microarchitecture of native ECM, which is known to support cell adhesion, signaling, and proliferation^49^. Although macroscopic analysis suggested uneven scaffold distribution across the implant surface (Figure 1F, G, I, J), EDS analysis demonstrated a consistent coating of organic matrix. Even in regions that macroscopically seem uncoated, the surface exhibited significantly elevated carbon content, indicating successful and uniform scaffold integration (Figure 1N, O, Table S.1).

### 2.2. *In vitro* Scaffold Integrity and HA Degradation

The stability of the scaffolds and the contributions of each of the components, FmocFF and HA were evaluated by comparing lyophilized hybrid scaffolds to implants integrated with each component separately (Figure 2). Hydrogels composed of FmocFF or HA alone, as well as the hybrid FmocFF/HA, were integrated into implants and subjected to lyophilization. Following rehydration in MEMα medium and incubation at 37° C, distinct differences in structural integrity were observed. Implants containing HA alone dissolved almost immediately upon rehydration, demonstrating the limited stability of HA in aqueous environments (Figure 2A, B, C). In contrast, pure lyophilized FmocFF scaffold remained intact during the first 24 hours of rehydration and began to disintegrate into the medium within seven days of incubation (Figure 2D, E, F). Notably, the hybrid FmocFF/HA scaffold maintained its structure without significant disintegration over seven days of incubation in the media (Figure 2G, H, I). These results suggest that integrating HA within the FmocFF network enhances scaffold stability, likely due to synergistic interactions that prevent rapid dissolution and improve structural integrity.

**Figure 2.**
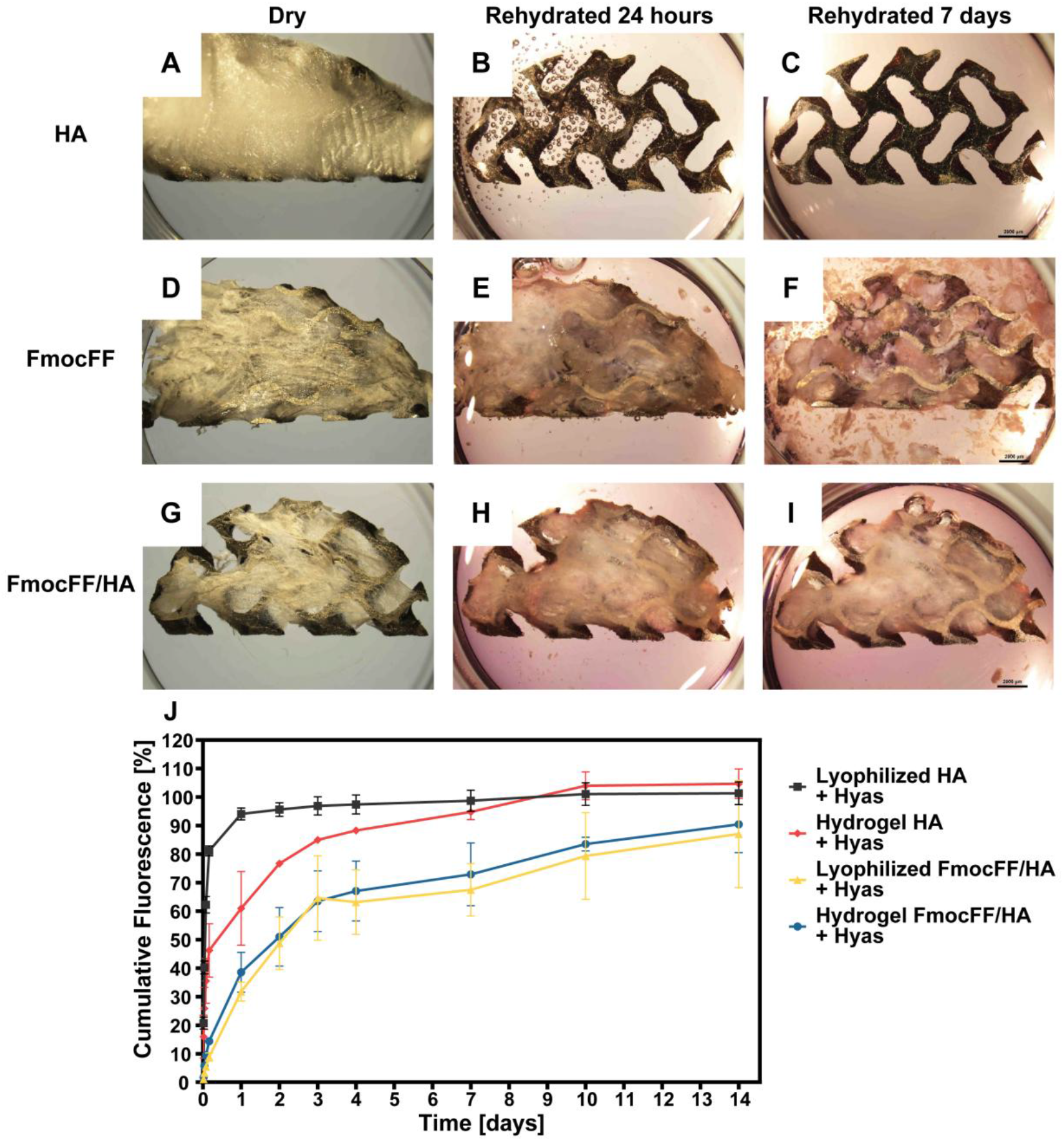
*In vitro* degradation dynamics of scaffold-integrated implants. (A–I) Media-induced rehydration of lyophilized scaffolds embedded within 3D-printed implants over 24 hours and 7 days at 37 °C. (A–C) HA-only lyophilized scaffold, (D–F) FmocFF-only lyophilized scaffold, and (G–I) composite FmocFF/HA lyophilized scaffold. (J) Cumulative degradation of fluorescently labeled HA in the presence of hyaluronidase (Hyas) when incorporated alone or as part of FmocFF/HA, in both hydrogel and lyophilized forms.

The enzymatic degradation profile of HA was assessed by incubating bioactive implants containing fluorescently labeled HA in hyaluronidase solution and monitoring fluorescence release over 14 days. Lyophilized HA alone degraded rapidly, with >80% of the HA lost within the first few hours. Although HA in solution degraded more slowly, it still exhibited significantly faster degradation than HA integrated into the FmocFF/HA hybrid scaffold, in either the hydrogel or the lyophilized form, which showed improved resistance to enzymatic cleavage (Figure 2J). These findings indicate that the hybrid scaffold effectively protects HA from rapid enzymatic degradation, highlighting its potential for enhanced stability in biomedical applications.

### 2.3. *In vitro* Cell-Scaffold Interaction

Next, we examined the *in vitro* cytocompatibility of the scaffold-integrated implants. The MG63 human osteosarcoma cell line, widely used as an osteoblast-like model, was employed to evaluate cell-scaffold interactions relevant to bone regeneration^67^. Cell attachment and morphology were assessed using HRSEM following 48 hours of MG63 osteoblast-like cell culture on inert implants, FmocFF/HA lyophilized-integrated implants, and hydrogel-integrated implants. In all groups, cells exhibited adhesion, spreading, and evidence of mitotic activity, indicating initial biocompatibility across the different surfaces (Figure 3).

**Figure 3.**
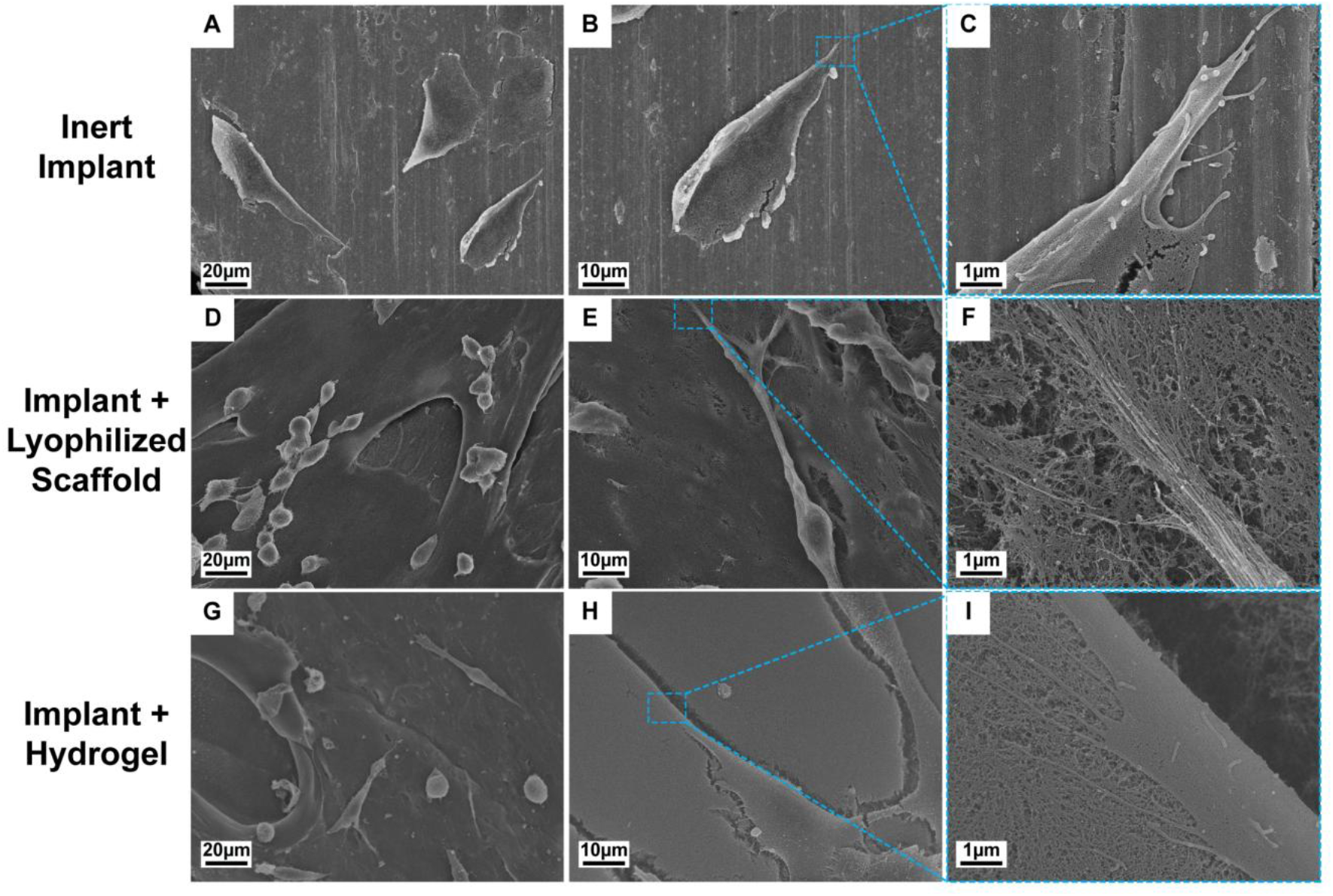
HRSEM analysis of cell-surface interactions on 3D-printed implants. (A-C) MG63 osteoblast-like cells cultured for 48 hours on the inert titanium implant. Sequentially higher magnification reveals the interface between the smooth titanium surface and adherent cells, with a clear distinction between cellular and substrate structures. (D-F) MG63 cells cultured on the FmocFF/HA lyophilized scaffold-incorporated implant. (G-I) MG63 cells cultured on the FmocFF/HA Hydrogel-incorporated Implant. At high magnifications, the bioactive scaffolds exhibit a nano-fibrillar architecture resembling native ECM, closely integrated with the attached cells. Scale bars: (A, D, G) 20 µm; (B, E, H) 10 µm; (C, F, I) 1 µm.

Although overall cellular morphology appeared similar, differences in cell-substrate interactions were evident. Unlike the inert implant (Figure 3A-C), on the hydrogel-integrated and lyophilized-integrated implants, cells were found to interact with a dense nanofibrillar network resembling the structural organization of the native ECM (Figure 3D–I). This fibrillary architecture, formed by the self-assembled FmocFF peptide network, likely provides essential biophysical and topographical cues that support cellular adhesion, cytoskeletal organization, and signaling pathways involved in adhesion and proliferation.

On the smooth, rigid surfaces of titanium implants, only limited focal adhesion points were typically observed. While titanium implant substrates provide essential mechanical support, they also influence cell behavior through mechanotransduction^47^. In contrast, within the nanofibrous environment of the FmocFF/HA scaffold, cells exhibited more abundant focal adhesions, likely due to the scaffold’s ability to recapitulate key features of the native extracellular matrix (ECM). This biomimetic architecture may facilitate more physiologically relevant cell–material interactions, resembling the *in vivo* preference of cells to anchor to fibrillar ECM proteins such as collagen, fibronectin, and laminin.

### 2.4. *In vivo* Critical-Size Rabbit Calvarial Defect for Restored Bone Regeneration

#### 2.4.1. Micro-CT assessment of the Entire Bone Defect

The regenerative performance of the bioactive implants, utilizing two different incorporation strategies, was evaluated in a rabbit calvarial critical-size defect model. Three defects 8 mm in diameter were created in the calvaria of nine female New Zealand White rabbits. Each defect was reconstructed with one of the following treatments: (i) an inert porous Ti-6Al-4V implant (control), (ii) an implant integrated with a lyophilized FmocFF/HA scaffold, or (iii) an implant integrated with FmocFF/HA hydrogel. All implants were fabricated by additive manufacturing using SLM to produce gyroid cylinders with a 5×5×5 mm cell size and a wall thickness of 0.3 mm (Figure 4A).

**Figure 4.**
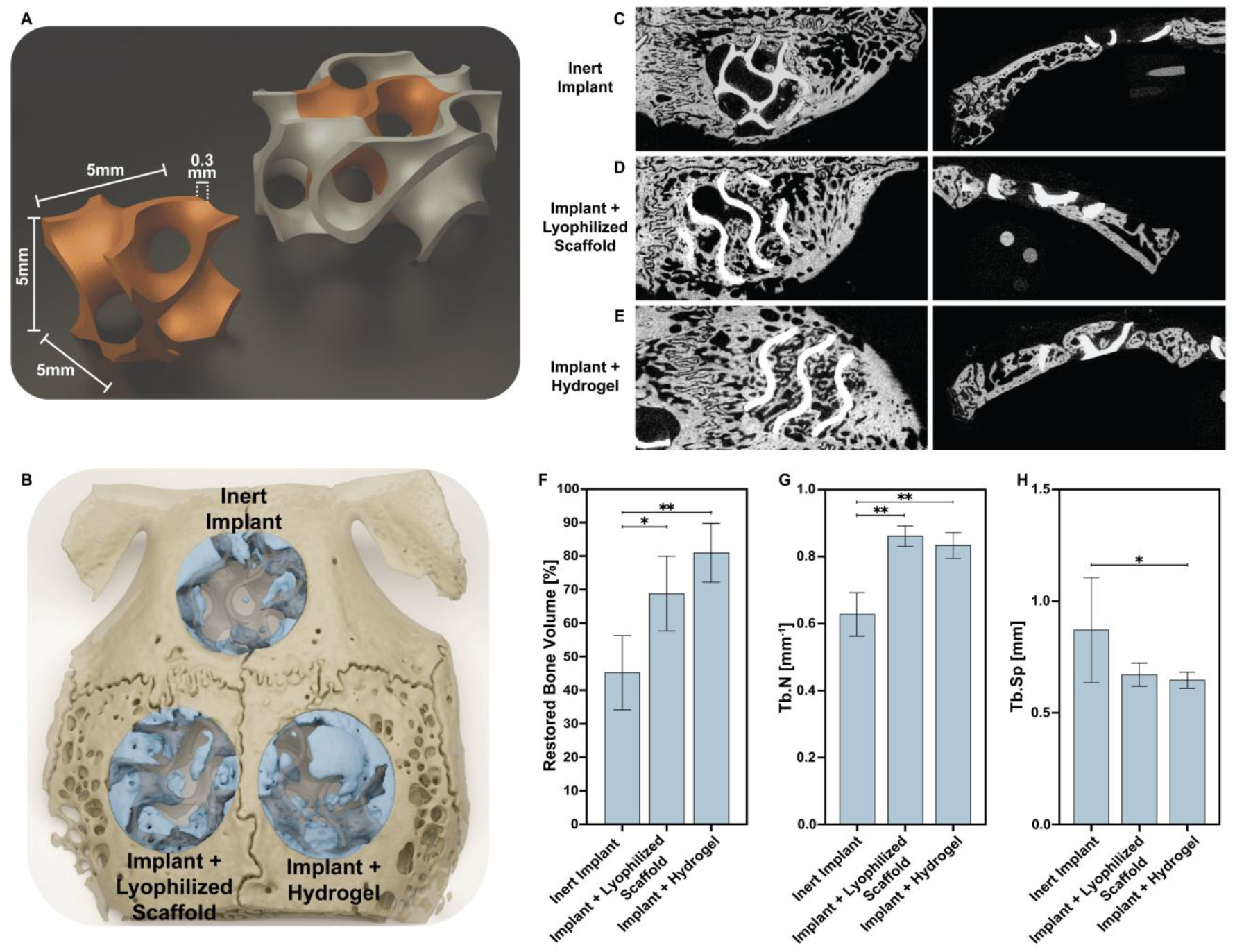
Micro-CT assessment of *in vivo* bone regeneration in rabbit calvarial defects treated with bioactive titanium implants. (A) Schematic illustration of the gyroid lattice unit cell used for implant fabrication, with 5×5×5 mm with cell size and 0.3 mm wall thickness. (B) 3D micro-CT reconstruction of a rabbit calvaria illustrating bone regeneration across the three implant types. Regenerated bone is shown in light blue, pristine bone in light brown, and implants in semi-transparent grey. (C-E) Representative micro-CT cross-sections (top and lateral views) of calvarial defects 8 weeks post-surgery in rabbits treated with: (C) inert titanium implant, (D) titanium implant incorporating lyophilized FmocFF/HA scaffold, and (E) titanium implant incorporating FmocFF/HA hydrogel. (F) Quantification of restored bone volume within the entire defect area for each treatment group. (G) Trabecular number calculation and (H) Trabecular separation measurements of the regenerated bone (mean ± SD; *p < 0.05, **p < 0.01, two-way ANOVA).

Animals were euthanized at 8 weeks post-surgery, and calvarial specimens were harvested for micro-CT and histological analyses. Visual assessment of the micro-CT sections indicated some degree of implant integration across all groups. However, substantially suprior bone formation was observed in the central regions of defects treated with the FmocFF/HA-integrated implants, both in hydrogel and lyophilized forms. In contrast, bone regeneration in the inert control group was predominantly limited to the periphery of the defect and regions adjacent to native bone. Notably, the hydrogel-integrated implants exhibited pronounced bone ingrowth extending throughout the inner zones of the implant architecture (Figure 4C-E and Figure S.2).

Quantitative micro-CT and 3D reconstruction analyses confirmed that incorporation of FmocFF/HA, whether as a hydrogel or as a lyophilized scaffold, significantly enhanced overall bone regeneration compared to the inert implants. Although the difference did not reach statistical significance, there was a trend toward higher bone volume in the hydrogel-integrated group (81%) compared to the lyophilized group (68.6%), both of which outperformed the inert implant group (48.4%) (Figure 4F). Morphometric analysis further demonstrated an increased trabecular number (Tb.N) in both FmocFF/HA-integrated groups, 0.84 mm^−1^ for the hydrogel group and 0.86 mm^−1^ for the lyophilized group, compared to 0.62 mm^−1^ in the inert control, indicating a denser trabecular architecture and improved bone quality for the scaffold-treated implants (Figure 4G). Moreover, trabecular separation (Tb.Sp) was notably higher in the inert implant group (0.88 mm) relative to the lyophilized (0.68 mm) and hydrogel (0.65 mm) groups, suggesting poorer bone regeneration and disrupted trabecular connectivity surrounding the inert implant (Figure 4H).

#### 2.4.2. Regional Analysis of Bone Regeneration

The central region of a bone defect is often recognized as particularly challenging for successful repair in critical-size defect models^8,14,15,20^. Thus, although titanium alloys such as TiAl6V4 are known to exhibit excellent osseointegration^10,12,45^, new bone formation often remains concentrated near the defect margins, leaving the central regions inadequately regenerated. This limited ingrowth can compromise long-term implant stability and hinder full integration with the surrounding bone^20^. To gain deeper insights into the spatial pattern of bone regeneration, the defect volume was virtually divided into two volume-equal segments, the outer and inner regions.

Analysis of bone formation within the inner and outer zones of the defect revealed a more pronounced regenerative effect of the hydrogel in the central region (Figure 5). In the inner zone, the hydrogel-integrated group exhibited nearly a twofold increase in bone volume (43.8%) compared to the inert control (21%). The lyophilized-integrated group also showed average enhanced bone formation in this region (32.3%), although the difference compared to the inert group did not reach statistical significance. In the outer zone, both FmocFF/HA-integrated groups improved bone regeneration, with bone volume reaching 106% in the lyophilized-integrated group and 101% in the hydrogel-integrated group, compared to only 78% in the inert control.

**Figure 5.**
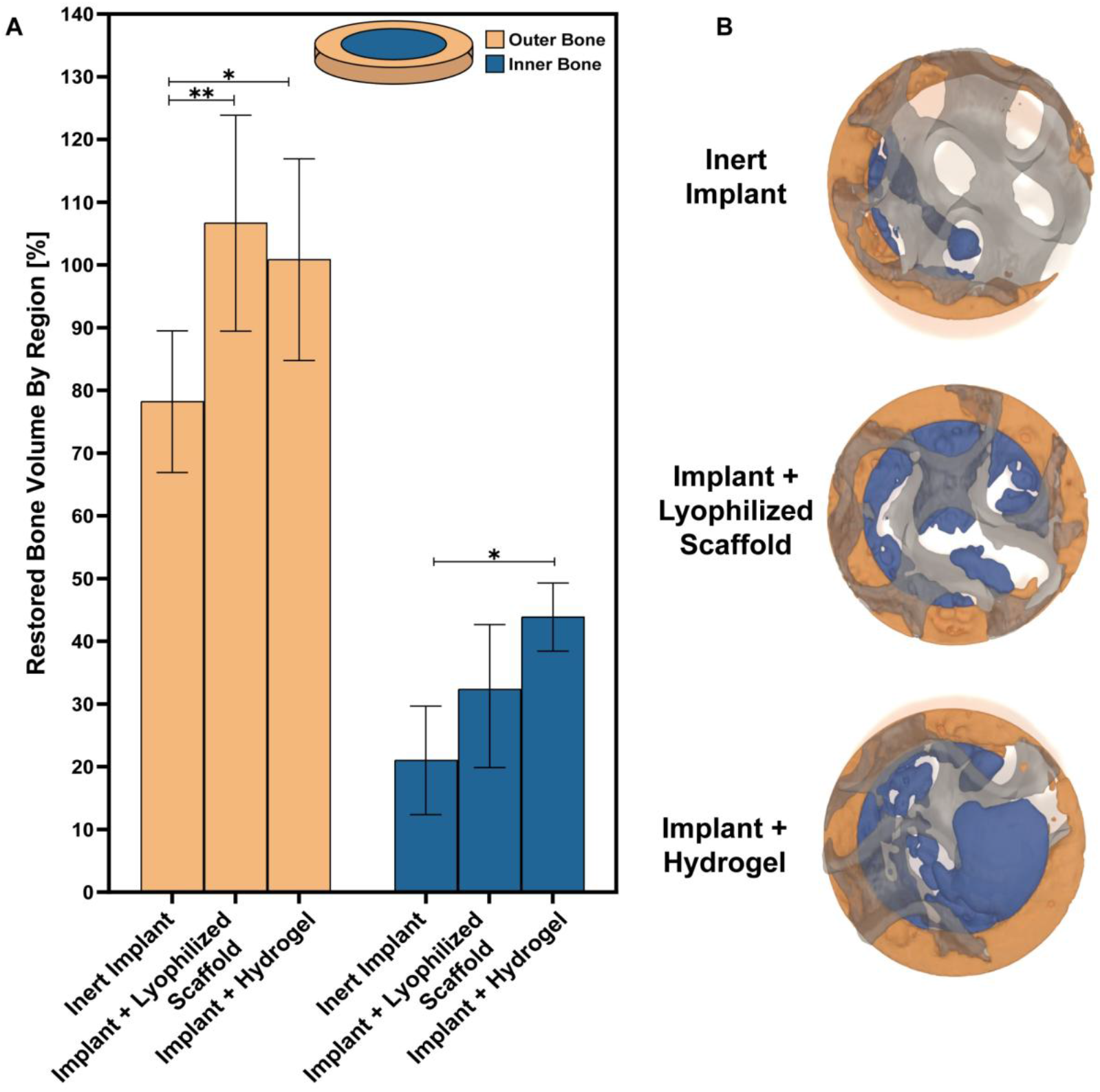
Regional micro-CT analysis of regenerated bone volume. (A) Quantification of regenerated bone volume within defined outer and inner regions of the calvarial defect, each occupying half of the total defect volume (two-way ANOVA: *p < 0.05, **p < 0.01). (B) Representative 3D micro-CT reconstructions of selected defects from each treatment group, showing digitally segmented regenerated bone in outer and inner regions. The color scheme corresponds to the group legend in the graph presented in (A).

These findings highlight the importance of the incorporation of bioactive scaffolds in achieving deep and uniform bone regeneration throughout the defect volume, potentially overcoming one of the key limitations of traditional metallic implants.

#### 2.4.3. Histological Analysis

Calvarial specimens were sectioned using a microtome (Figure 6A-C) and subjected to hematoxylin and eosin (H&E) (Figure 6D-I) and Masson’s Goldner trichrome (MGT) (Figure 6J-O) staining. The histological findings were consistent with the micro-CT results, demonstrating the contribution of FmocFF/HA, either in its lyophilized form or as a hydrogel, to the promotion of osteogenesis and bone ingrowth toward the implant core.

**Figure 6.**
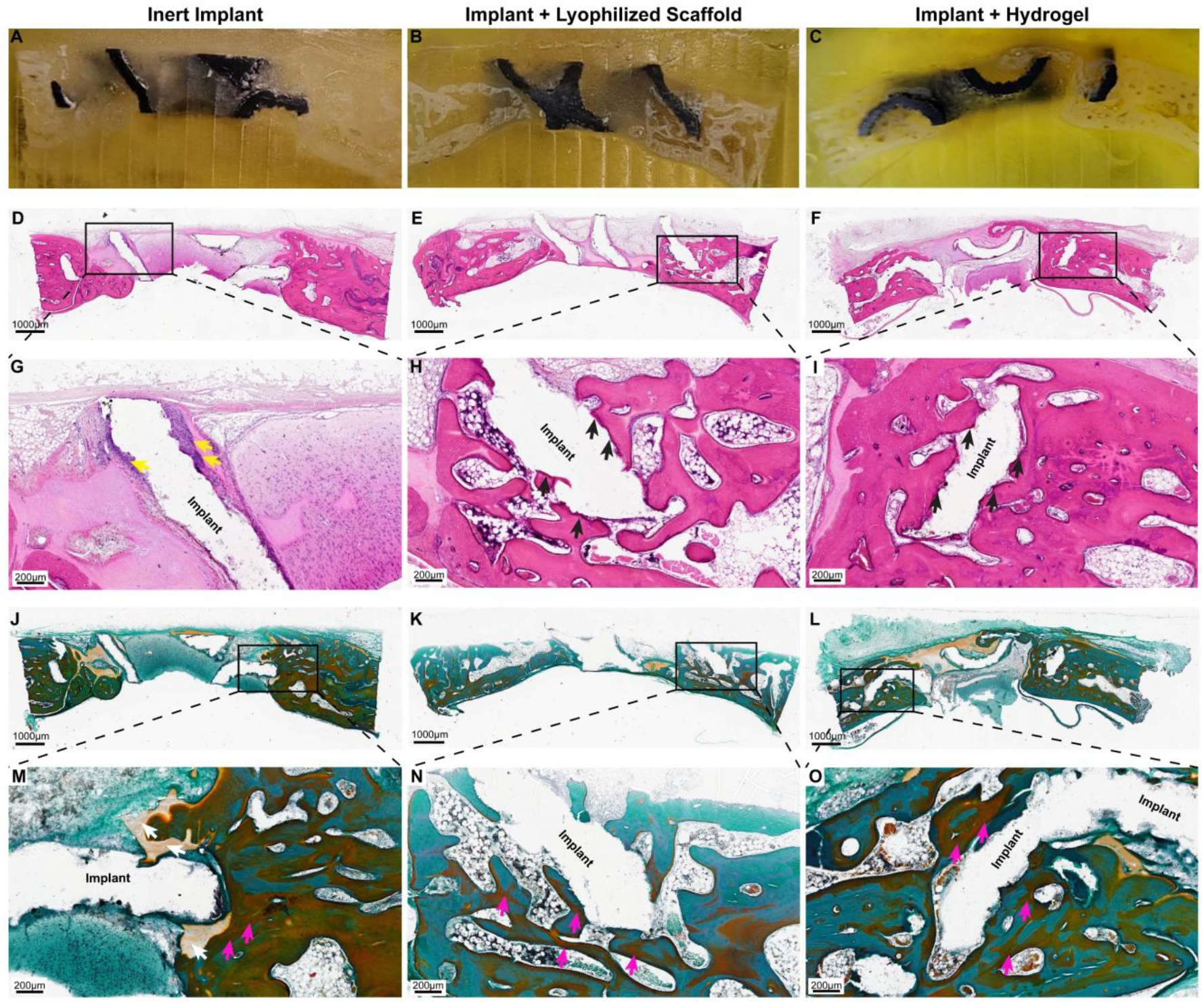
Histological analysis of the regenerated bone. (A–C) Overview images of undecalcified bone sections during preparation, prior to laser microtome removal of metallic components. (D–I) H&E staining of tissues adjacent to the implants. (D–F) Low-magnification images of entire sections; (G–I) Higher magnification views of selected regions of interest (ROIs). In H&E staining, bone tissue appears dark pink (black arrows), whereas connective tissue exhibits a lighter pink coloration. Inflammatory areas appears purple (yellow arrows). The area previously occupied by the implant appears as a white void, indicated in the magnified images. (J–O) MGT staining to evaluate bone formation, osteoid presence (magenta arrows), and active periosteum. (J– L) Low-magnification images of entire sections; (M–O) Higher magnification images of specific ROIs. In this staining, mature bone appears bright to dark turquoise, osteoid is brown-orange, dense connective tissue is bright orange, loose connective tissue appears blue, dense connective tissue appears bright orange (white arrows) bone marrow stains dark purple, and active periosteum is orange. The former implant space is visualized as a white void and labeled in the enlarged images. Images correspond to the following experimental groups: (A, D, G, J, M) inert implant; (B, E, H, K, N) lyophilized FmocFF/HA-incorporated implant; (C, F, I, L, O) implant combined with FmocFF/HA hydrogel. (D-F, J-L) Scale bars are 1000µm, (G-I, M-O) Scale bars are 200µm.

In the inert implant group, H&E staining revealed moderate inflammatory infiltrate surrounding the implant (Figure 6G, yellow arrows), suggesting a mild foreign body response. Notably, this inflammatory reaction was absent in both FmocFF/HA-integrated groups, indicating a potentially beneficial modulatory effect of the bioactive components on local tissue responses (Figure 6H, I).

In both the lyophilized and hydrogel FmocFF/HA groups, the bone marrow appeared healthy and active, even in regions immediately adjacent to the implant surface. Furthermore, significant new bone formation was observed (Figure 6H, I, black arrows), accompanied by pronounced active periosteum and osteoid formation in both bioactive groups (Figure 6N, O, magenta arrows). These findings suggest that FmocFF/HA not only supports osteoconduction but may also stimulate osteoinductive processes, enhancing bone remodeling activity near the implant. The hydrogel group exhibited a higher level of direct bone-implant interaction, as evidenced by more continuous bone tissue in close proximity to the lattice structure (Figure 6I, black arrows). This may reflect on an improved diffusion and spatial distribution of the hydrogel within the implant’s porous architecture.

Although the inert implant did not actively promote bone formation, its bioinert properties allowed for some degree of bone integration surrounded by dense connective tissue at the periphery of the implant (Figure 6M, white arrows). This limited peripheral integration underscores the importance of bioactive modifications, such as FmocFF/HA incorporation, in achieving substantial bone ingrowth and osseointegration throughout the implant structure.

## 3. Conclusion

This study presents the successful integration of bioactive FmocFF/HA hybrid hydrogel and its lyophilized scaffold into 3D-printed porous Ti-6Al-4V implants, enhancing bone regeneration in critical-size defects in a rabbit calvaria model over eight weeks. The nanofibrillar scaffold mimics the ECM, improving cell adhesion and enzymatic stability compared to HA alone. *In vivo* results showed that the integration of both hydrogel and lyophilized scaffolds into the implants significantly increased bone formation compared to inert implants throughout the defect, with the hydrogel achieving the highest regeneration. Notably, the amount of newly formed bone in the inner region of the implant was doubled in the hydrogel group compared to the inert group. This work presents a novel, simple, cell-free strategy, without any addition of growth factors or cytokines, for fabricating bioactive implants for personalized bone reconstruction, opening new avenues for next-generation therapies in regenerative medicine.

## 4. Materials and Methods

### 4.1. Materials

Fmoc-FF was purchased from GL Biochem Ltd. (Shanghai, China). LaserForm Ti Gr23 (A) powder was obtained from 3D Systems (Rock Hill, SC, USA), and 3D-printed implants were supplied by Sharon Tuvia (1982) Ltd. (Ness Ziona, Israel). Hyaluronidase enzyme powder, dimethyl sulfoxide (DMSO), and fluorescein-conjugated hyaluronic acid (Fluorescein-HA) were acquired from Sigma-Aldrich (Rehovot, Israel). Pre-filled syringes containing 1% (w/v) HA in phosphate-buffered saline (PBS) were purchased from BTG-Ferring (Kiryat Malachi, Israel). Minimum Essential Medium Alpha (MEM-α), fetal bovine serum (FBS), penicillin–streptomycin solution, and non-essential amino acids were obtained from Biological Industries (Beit HaEmek, Israel).

### 4.2. Ti-6Al-4V Lattice Implant Design and Fabrication

The lattice implants were designed using 3DXpert software (3D Systems, Rock Hill, South Carolina, USA) and fabricated from Ti-6Al-4V powder (LaserForm Ti Gr23, 3D Systems) by selective laser melting (SLM) using Flex 350 printer (3D Systems, Rock Hill, South Carolina) at Sharon Tuvia (1982) Ltd. (Ness Ziona, Israel). Two cylindrical structures 50 mm in length and 8 mm in diameter with a gyroid lattice architecture (unit cell size of 5×5×5 mm) were printed. Post-processing included standard heat treatment for stress relief, mechanical removal of supports, and dry electropolishing (DLyte, DryLyte Technology, Barcelona, Spain). Subsequently, the cylinder was sectioned into discs 1.8 mm in thickness using an IsoMet® 1000 Precision Diamond Saw (Buehler, Chicago, USA). The implants were cleaned and sterilized by autoclaving at 135 °C prior to incorporation of the bioactive component, which was performed under sterile conditions. In addition, for *in vitro* experiments aimed at assessing the penetration of the bioactive material into the core of the implant, cylindrical implants 10 mm in height and 15 mm in diameter were fabricated. These implants were designed to be easily split into two semi-cylindrical halves, enabling visual and microscopic evaluation of material distribution throughout the implant’s core. Fabrication and post-processing of these implants followed the same SLM parameters and procedures described above.

### 4.3. Integration of the Peptide-Polysaccharide Hydrogel

The FmocFF/HA hydrogel was prepared using a solvent-switch method, as described previously^64,66^. Briefly, lyophilized FmocFF powder GL Biochem Ltd. (Shanghai, China) was dissolved in DMSO at a concentration of 100 mg/mL. HA (BTG Ltd. Israel) was initially supplied as a 1% (w/v) stock solution and diluted with double-distilled water (DDW) to a final concentration of 0.167% (w/v). The diluted HA solution was mixed on an orbital shaker for 2 hours to ensure complete dissolution. The implants were immersed in the diluted HA solution prior to the addition of the peptide solution. To 1 mL of the prepared HA solution, 50 µL of the FmocFF/DMSO stock was added, yielding final concentrations of 0.5% (w/v) FmocFF and 0.167% (w/v) HA. The mixture was briefly vortexed to promote uniform dispersion and then incubated at 4 °C for at least 4 hours to allow self-assembly. For lyophilized samples, hydrogel-embedded implants were first frozen at -20 °C for 1 hour, then transferred to -80 °C for deep freezing. Implants were subsequently lyophilized overnight while still embedded in the hydrogel matrix, resulting in constructs uniformly filled and coated with lyophilized peptide-polysaccharide gel.

### 4.4. High Resolution Scanning Electron Microscopy (HRSEM)

Implants integrated with FmocFF/HA in either hydrogel or lyophilized form were prepared as described in Section 2.3. Following complete gelation, hydrogel samples were fixed in 2.5% (v/v) glutaraldehyde at 4 °C overnight. The next day, samples were dehydrated through a graded ethanol series (25%, 50%, 75%, 95%, and 100%), with each step lasting 15 minutes. The samples were then coated with an 8 nm layer of gold (Au) to enhance surface conductivity.

High-resolution imaging was performed using a Zeiss GeminiSEM 300 scanning electron microscope (Carl Zeiss, Germany). Elemental analysis of selected regions was conducted using energy-dispersive X-ray spectroscopy (EDS) integrated into the HRSEM system.

### 4.5. Lyophilized Scaffold Rehydration

HA, FmocFF solution, and FmocFF/HA were incorporated into the implants as outlined in section 2.3, followed by lyophilization. Images of the dry scaffold-incorporated implants were taken, and then 2 mL of 10% FBS MEM-Alpha medium was added to each of the samples. The samples were incubated inside a cell culture incubator at 37 °C and 5% CO_2_. Images were taken after 24 hours and 7 days of incubation, using a zoom stereomicroscope (Nikon SMZ800N) equipped with a DS-Fi2 camera (Nikon, Japan).

### 4.6. *In vitro* Enzymatic Degradation by Hyaluronidase

Bovine testicular hyaluronidase (Type IV-S, lyophilized powder; Sigma-Aldrich, Rehovot, Israel) was reconstituted in 100 mM sodium acetate buffer supplemented with 2 mM CaCl_2_. To enable fluorescence-based monitoring of HA degradation, Fluorescein-HA (Sigma-Aldrich, Rehovot, Israel) was incorporated into the HA solution at a concentration consistent with that of the original unlabeled HA. Lyophilized FmocFF/HA-integrated implants and FmocFF/HA hydrogel-integrated implants were incubated in enzyme solution containing 200 U/mL hyaluronidase at 37 °C to evaluate enzymatic degradation kinetics. All samples were maintained in the dark throughout the experiment to minimize photobleaching of the fluorescent probe. At predefined time intervals, the enzyme solution was carefully removed and replaced with fresh aliquots to ensure sustained enzymatic activity. The collected supernatants were analyzed for fluorescence intensity at 485 nm excitation and 535 nm emission using a TECAN Spark microplate reader (Tecan, Switzerland). Implants containing pure HA hydrogels served as positive controls for enzymatic degradation.

### 4.7. Examination of the Cell-Scaffold Interaction

Human osteosarcoma MG-63 cells (ATCC - CRL-1427 ^™^) were cultured in MEM-α medium supplemented with 10% FBS, 1% penicillin–streptomycin, and 1% non-essential amino acids. Inert, lyophilized-integrated, and hydrogel-integrated implants were prepared as described above. Each implant was placed in a well of a 12-well cell-repellent culture plate, and 2 × 10^5^ cells were seeded onto each sample.

After 48 hours of incubation, the culture medium was removed, and the samples were washed twice with PBS, followed by overnight fixation in 2.5% glutaraldehyde at 4 °C. The next day, samples were washed twice with PBS and dehydrated through a graded ethanol series (25%, 50%, 75%, 95%, and 100%; 10 minutes at each step). Critical point drying was then performed. For imaging, samples were sputter-coated with an 8 nm gold layer and analyzed using a Zeiss GeminiSEM 300 high-resolution scanning electron microscope (Carl Zeiss, Germany).

### 4.8. *In vivo* Rabbit Calvarial Critical Size Bone Defect Model

Adult female New Zealand White rabbits (2.7-3.3 kg) were subjected to calvarial bone defects. The experimental procedures involving live vertebrates in this study were approved by the institutional committee of animal care and use (IACUC) of Tel Aviv University (no. TAU - MD - IL - 2306 - 143 - 4), and all experiments were strictly conducted in accordance with the approved guidelines. Animals were individually housed in the Central Animal Facility of Tel Aviv University. Animals were fed with Teklad Global Rabbit Diet (Envigo, Madison, Wisconsin), autoclaved hay, and tap water *ad libitum*.

Surgical procedure was performed under general anesthesia following sedation with 5 mg/kg subcutaneous (SC) xylazine (Sedaxylan Veterinary, Eurovet Animal Health BV Bladel, The Netherlands) and 35 mg/kg SC ketamine (Clorketam, Vetoquinol, Lure, France). Oxygen was delivered by means of a facemask connected to an anesthetic machine with an oxygen flowmeter (model Mix4; Foures SAS, Bordeux) at a rate of 2 l/min. Blood oxygen saturation, pulse, and body temperature were monitored using a PhysioSuite monitor (Kent Scientific). Once anesthetized, the surgical area was shaved and disinfected with iodine solution, and local anesthesia was administered with 2% lidocaine hydrochloride and norepinephrine (1:100,000) for the reduction of pain and hemostasis. The rabbit calvarium was exposed via a U-shaped incision. The periosteum was carefully separated from the bone and skin flaps were elevated to expose the calvarium. Three circular defects 8 mm in diameter were made in the bone under saline irrigation. To mark the dimensions of the defects, an 8-mm trephine bur was used, followed by a gentle bone removal process using an inverted conical diamond bur. Attention was given to avoid damaging the dura mater. The three treatment modalities were randomly allocated for all 27 defects, with three defects per animal: 1. Inert 3D-printed implant. 2. FmocFF/HA “wet” hydrogel-integrated 3D-printed implant. 3. FmocFF/HA “dry” lyophilized-integrated 3D-printed implant. To assess the non-unity of the defect, four defects remained unfilled. The conditioned implants were prepared as described in Section 2.3. Representative images of the surgical procedure workflow are shown in figure S.1. Eight weeks after the implantation, the rabbits were euthanized in accordance with ethical guidelines. The calvaria from each rabbit was harvested and fixed in 10% neutral buffered formalin (Sigma-Aldrich, Rehovot, Israel) for subsequent micro-computed tomography (micro-CT) and histological analyses.

### 4.9. Micro-CT Analysis of the Bone and the Implants

All calvaria were scanned using Micro-CT (XT H 225 ST, Nikon Metrology NV, Leuven, Belgium), equipped with a 225 kV 225 W reflection target. Scans were performed at an isotropic resolution of 25 μm utilizing the following parameters: 180 kVp energy, at 133 μA intensity, a 0.5 mm Tin filter, with 1570 projections utilizing a 354 msec exposure time. Raw scans underwent reconstruction procedure using the Nikon CT Pro 3D software (v. 6.9.1; Nikon Metrology NV, Leuven, Belgium) and subsequent segmentation using various semiautomatic tools based on grayscale thresholds with appropriate manual refinement prior to the analysis in Amira software (v. 6.3, www.fei.com). Each scan was segmented into bone and implant, with the latter clearly visible and morphologically distinguishable from the surrounding bone. Circular regions of interest (ROI) 8 mm in diameter were identified via the surgical borders and subjected to volumetric analysis. Quantitative morphometric analysis included calculation of the trabecular number and trabecular separation within defined regions. Each 8 mm circular defect was virtually divided into two regions (outer and inner) using two coaxial 3D cylinders of varying radii (modelled to divide the defect area into two identical volumes). The circular ROI was subjected to adjacent intact bones to serve as a reference of a normal bone. Volumes of the restored bone were calculated as a ratio of the bone volume present in each and to the control volume of the adjacent pristine bone that served as control.

### 4.10. Histological Processing and Analysis of the Bone

Histological processing was carried out at Patho-Logica Laboratories (Ness Ziona, Israel). Samples were dehydrated and embedded in Spurr epoxy resin for non-decalcified sectioning. Plastic blocks were trimmed and sectioned using a laser microtome (TissueSurgeon, LLS ROWIAK LaserLabSolutions, Hannover, Germany), employing a femtosecond near-infrared laser for precise, non-contact cutting of implant-containing hard tissues with minimal material loss. Serial transverse sections, 12 μm thick, were prepared, mounted on glass slides, and stained using Hematoxylin and Eosin (H&E) for general morphology and with Masson-Goldner Trichrome (MGT) for connective tissue visualization. Slides were scanned using a KF-BIO-40 slide scanner.

### 4.11. Statistical Analysis

Statistical analyses were performed using GraphPad Prism (version 10.4.1, GraphPad Software, San Diego, CA, USA). For comparisons involving more than two groups, one-way or two-way analysis of variance (ANOVA) was used as appropriate, followed by Tukey’s post hoc test to assess differences between individual groups. Statistical significance was defined as *p* < 0.05. All quantitative data are presented as mean ± standard deviation (SD). Significance levels are indicated in the figures as follows: \**p* < 0.05, **p < 0.01, and corresponding group comparisons are annotated with symbols as defined in the respective figure legends.

## Author Information

## Acknowledgment

Noam Rattner thanks the Sagol Center for Regenerative Medicine for financial support. The authors acknowledge the staff at the Gray Faculty of Medical and Health Sciences Research Infrastructure Core Facilities of Tel-Aviv University and the Chaoul Center for Nanoscale Systems of Tel Aviv University for the use of instruments and staff assistance. We thank the Dan David Center for Human Evolution and Biohistory Research for the use of instruments and staff assistance. We are grateful to Sharon-Tuvia Ltd. for the custom manufacturing of the 3D-printed implants and Patho-Logica Ltd. for histological processing assistance. We thank the Adler-Abramovich group and Levin Center members for helpful discussions.

## Data Availability Statement

The data that support the findings of this study are available from the corresponding author upon reasonable request.

